# Fine-tuning of deep language models as a computational framework of modeling listeners’ perspective during language comprehension

**DOI:** 10.1101/2021.11.22.469596

**Authors:** Refael Tikochinski, Ariel Goldstein, Yaara Yeshurun, Uri Hasson, Roi Reichart

## Abstract

Computational Deep Language Models (DLMs) have been shown to be effective in predicting neural responses during natural language processing. This study introduces a novel computational framework, based on the concept of fine-tuning (Hinton, 2007), for modeling differences in interpretation of narratives based on the listeners’ perspective (i.e. their prior knowledge, thoughts, and beliefs). We draw on an fMRI experiment conducted by Yeshurun et al. (2017), in which two groups of listeners were listening to the same narrative but with two different perspectives (cheating versus paranoia). We collected a dedicated dataset of ~3000 stories, and used it to create two modified (fine-tuned) versions of a pre-trained DLM, each representing the perspective of a different group of listeners. Information extracted from each of the two fine-tuned models was better fitted with neural responses of the corresponding group of listeners. Furthermore, we show that the degree of difference between the listeners’ interpretation of the story - as measured both neurally and behaviorally - can be approximated using the distances between the representations of the story extracted from these two fine-tuned models. These models-brain associations were expressed in many language-related brain areas, as well as in several higher-order areas related to the default-mode and the mentalizing networks, therefore implying that computational fine-tuning reliably captures relevant aspects of human language comprehension across different levels of cognitive processing.

## Introduction

The meaning of each word in natural language is defined by its use in context. For example, the meaning of the word “cold” varies in the context of cold weather, cold personality, and cold symptoms. Typically, context is detriment by the relation of each incoming word to the preceding words in the text, and such dependencies can span multiple timescales: from the relationship among adjacent words, up to long-range dependencies across paragraphs. In agreement with such behavioral observations, studies using naturalistic stimuli have identified a topographical hierarchy of processing timescales extending from the early sensory cortex to higher-order areas (Hasson et al., 2008; Van Berkum, 2008; Lerner et al., 2011; Hasson et al., 2015). The responses in early sensory areas, such as auditory cortices, are influenced by the temporal structure over tens of milliseconds. These findings are consistent with research suggesting that early sensory areas process fast-changing, low-level stimulus features, such as phonological information. By contrast, neural responses in midlevel areas, which include linguistic regions along the superior temporal gyrus, are affected by information integrated over a few seconds (for example, the preceding words in the sentence). At the apex of the processing hierarchy, the neural response of the Default Mode Network (DMN) to each sentence is influenced by prior information accumulated over many minutes (Yeshurun et al., 2021; Friederici, 2020).

The realization that the meaning of each word in natural language is defined by its use in context was utilized recently to build deep language models (DLMs). DLMs (like BERT or GPT2) are trained to predict the appearance of words in text given the context (the other words in the text). In contrast to traditional symbolic psycholinguistic models, DLMs learn language from real-world textual examples “in the wild,” i.e., with minimal or no explicit prior knowledge about the structure or other linguistic properties of the input text. Furthermore, DLMs learn to encode a sequence of words into numerical vectors, termed contextualized embeddings, from which the model decodes the missing words. Interestingly, recent studies have demonstrated that contextual embeddings derived from DLMs (such as BERT or GPT2) can be used to predict neural responses in brain language areas during the processing of human language (Jain & Huth, 2018; Schwartz et al., 2019; Goldstein et al., 2021). Furthermore, a recent study used a DLM to model the topography of timescales of language processing, by varying the contextual window of the model, from a short contextual window of a few words up to a large contextual window that aggregates information across many paragraphs (Caucheteux et al., 2021). The similarities between deep language models and the way the brain processes language provide a new unified modeling framework for the study of the neural basis of the human language faculty.

Context, however, is not defined solely by the other words in the text, but also by the listeners’ perspective, as defined by their thoughts, beliefs, and knowledge. In such cases, the exact same words in a story, essay, or political speech, can have substantially different meanings across different readers with different backgrounds. An fMRI experiment that demonstrates this aspect of contextual information was conducted by Yeshurun et al. (2017). In their experiment, two groups of participants listened to the same audio recording of a short story by J.D. Salinger (‘Pretty Mouth and Green My Eyes’). In the story, a husband loses track of his wife at a party and returns alone to their apartment in the city. Worried and anxious he calls his best friend, in the middle of the night, about the whereabouts of his wife. Next to the best friend, in bed, lies a mysterious woman, whose identity is kept intentionally vague. Is she the wife, having an affair with the best friend (cheating context), or is she the friend’s girlfriend and the husband is unreasonably jealous as his friend implies (paranoia context)? Deciding among these two contexts will have great consequences for the interpretation of the conversation. Before listening to the story, each experimental group was primed to adopt only one of these interpretations. Listener’s perspective (cheating vs. paranoia) affected the neural responses to the story in areas with a long processing timescale, including the default mode network (DMN; Mars et al., 2012, Yeshurun et al., 2021), and frontal areas related to high-level language processing (Adolphs, 2009; Fletcher et al., 1995; Mar 2011).

Can we use DLMs as a computational framework to model how context is shaped by the listeners’ perspective (state of mind)? In this paper, we propose to use the concept of “fine-tuning” (Hinton, 2007) to model such an effect. DLMs are typically pre-trained on very large textual corpora (billions of words) sampled from a variety of textual domains and sources. This pre-training stage allows the model to learn how language is used across many natural contexts. Adapting the pre-trained model to perform a specific ‘downstream’ task (e.g., sentiment analysis), however, requires a second stage of training, known as the “fine-tuning” stage, in which the model’s parameters are subtly adjusted so that the language representations will better fit to the narrow context. In other words, the fine-tuning procedure dynamically reweights the DLM parameters that produce the textual representations (embeddings) to better fit the semantic space of a specific domain.

By this logic, we propose harnessing the fine-tuning technique in order to create different variants of a given DLM, each simulating human language processing through a lens of a different “state of mind”. To examine this idea, we fine-tuned the BERT language model (Devlin et al., 2018) to simulate two specific perspectives: paranoia and cheating. Our main aim is to test how fine-tuning the model’s context will improve our ability to model the listeners’ neural responses as they process Salinger’s story, using two very distinct states of mind.

## Method

### fMRI data

#### participants, stimuli, and experimental design

The current study reanalyzes a previously published fMRI dataset (Yeshurun et al., 2017). The dataset consists of fMRI scans of 40 right-handed subjects assigned to one of the following experimental conditions: *Cheating* (10 females, 10 males, age: *M*=20.85, *SD*=3.73) or *Paranoia* (9 females, 11 males, age: *M*=21.45, *SD*=3.42).

The stimulus was a 11 min and 32 seconds record of a professional actor reading a short story of J.D. Salinger: “Pretty Mouth and Green My Eyes.” The story describes a phone conversation between two friends, Arthur and Lee. Arthur has returned home after a party, and he lost track of his wife, Joanie. He is calling Lee to share his concerns over her whereabouts. Lee is at home, and a woman is lying on the bed next to him. The woman’s identity is ambiguous—she may or may not be Joanie, Arthur’s wife. Before listening to the story, participants were provided with a short introduction (~ 30 s) either specified that Arthur’s wife is cheating on him with Lee (for the cheating condition), or that Arthur is paranoid and that his wife is not cheating on him (for the paranoia condition; Figure 1). A story-comprehension questionnaire was administered immediately after the scan, and statistical analyses of the responses indicate that the context manipulation did affect the subject’s interpretation of the story (see Yeshurun et al., 2017 for more details).

**Figure 1.**
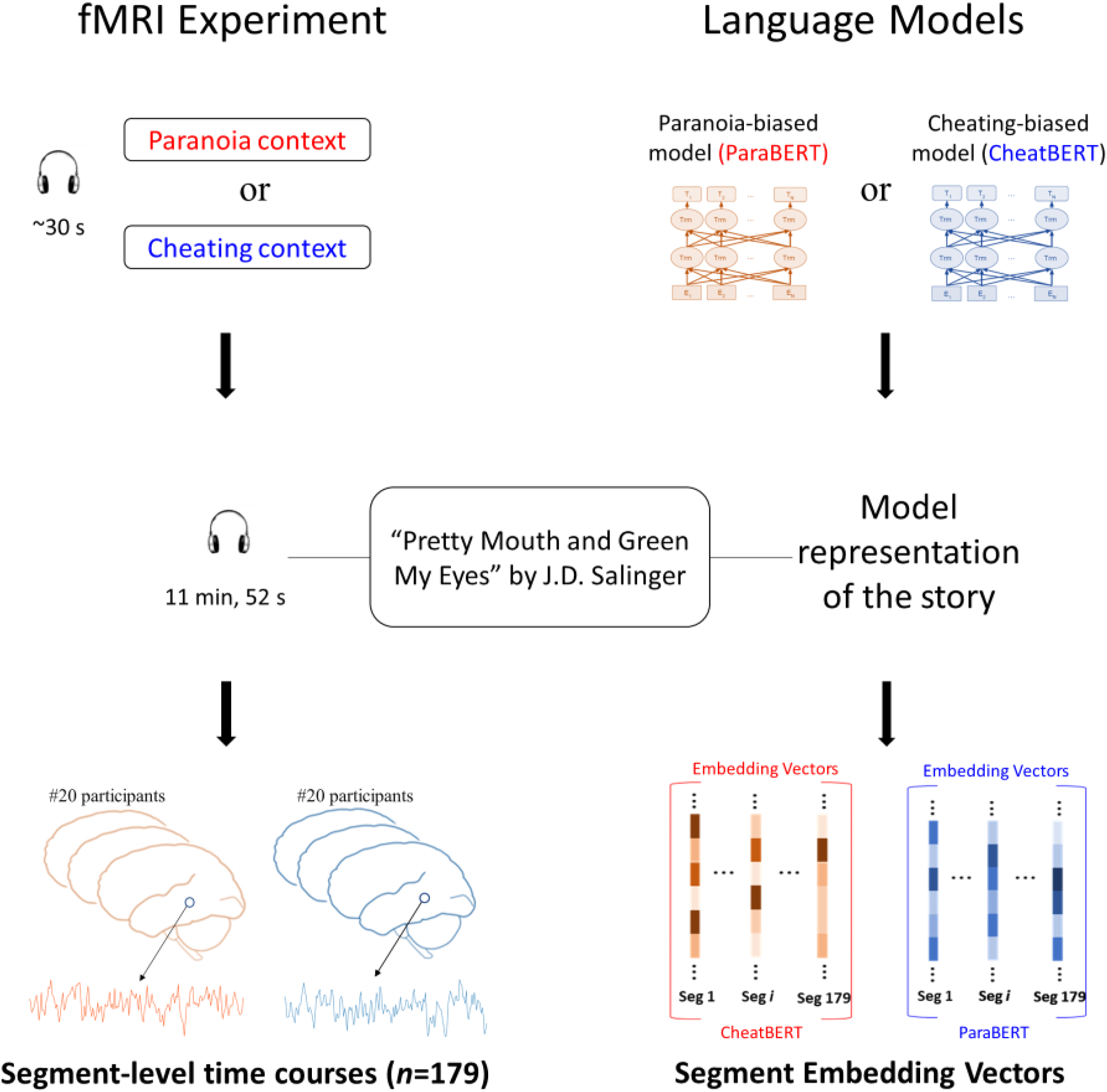
An illustration of both the neural and the computational context-dependent representations of the same narrative. The left side represents the fMRI experiment by which the neural representations were acquired: 40 subjects were listening to an ambiguous short story which can be interpreted in two main contexts, cheating or paranoia. Half of the subjects were primed to the cheating context, and the other half were primed to the paranoia context. The right side illustrates the computational modeling of the experiment. Two context-dependent language models were created, each simulates a different context. We used each model to extract vector representations of the story (a.k.a. embedding vectors).

#### Preprocessing and voxel selection

The functional MRI (fMRI) data were preprocessed by Yeshurun et al. (2017) and included the following steps: Motion correction, slice-time correction, linear-trend removal, high-pass filtering (two cycles per condition), spatial smoothing (Gaussian filter of 6-mm FWHM), spatial transformation to 3-D Talairach space (Talairach & Tournoux, 1988), and hemodynamic delay correction (based on the correlation between the audio envelope and the BOLD signal recorded from the A1+).

To filter out stimulus-irrelevant voxels, we executed a voxel-wise inter-subject correlation analysis (ISC, Hasson et al., 2004) across the whole gray matter: For each voxel, we isolated each subject’s time-course and correlated it with the averaged time-course of the remaining subjects. The voxel’s ISC-score is calculated by averaging the correlation scores (after Fisher’s Z transformation) obtained from repeating this process for all subjects. To assess how significantly each ISC-score is different from zero, we conducted a non-parametric permutation test by randomizing the phase of the signal (Simony et al., 2016) 1000 times prior to ISC calculation and used the obtained null distribution to estimate the p-value. We ran this procedure separately for each experimental group (cheating/paranoia) and selected for subsequent analyses only voxels that achieved a significant ISC-score (*p*<0.01, corrected for multiple tests using FDR) in both groups. The process yielded a total of 11783 ‘stimulus-locked’ voxels (Figure S2).

### Behavioral data

Besides neuroimaging data, we also used behavioral data collected by Yeshurun et al. (2017), which quantifies the effect of context on participant’s interpretation across the story. The text was divided into 179 segments (*Mean duration*=3.77 s, *SD*= 2.39 s) by an independent expert annotator, and five independent raters were asked to rate how differently participants from different groups (cheating/paranoia) would interpret each segment. The raw scores (on a scale from 1 to 5) were first standardized to Z-scores for each rater, and then averaged across raters. The inter-rater reliability was high, as reflected by a Cronbach’s α coefficient of 0.84 (See Yeshurun et al. (2017) for more details).

### Computational modeling

We propose a novel method for computational modeling of context modulation in story interpretation. The main idea is to take a pre-trained language model and modify (fine-tune) it toward either the cheating or the paranoia contexts. Specifically, we used a well-accepted language model – BERT (Devlin et al., 2019) – as our initial, context-independent language model, and designed a fine-tuning process that creates two new context-dependent variants of BERT – CheatBERT (for the cheating context) and ParaBERT (for the paranoia context). We administered the fine-tuning process using dedicated datasets and classification tasks as described below.

#### Fine-tuning tasks and datasets

We defined two binary classification tasks: One to distinguish between cheating stories and no-cheating stories, and the other to distinguish between paranoia stories and no-paranoia stories. Our basic hypothesis is that fine-tuning BERT on these tasks (i.e., updating its parameters, which were set in the pre-training stage) would bias its internal representation of language toward the cheating (when using the first fine-tuning task) context or the paranoia context (when using the second fine-tuning task).

For these tasks, we collected 2829 short stories (between 100 to 4096 words; average number of words= 757.27, *SD*=677.58) concerning matters of relationships and romance. The stories were written by users of the Medium.com of the Reddit.com websites. To locate relevant stories from Medium.com, the following website-tags were used: *marriage, relationships, romance, affairs, jealousy, monogamy, polygamy,* and *dating*. The stories from Reddit.com were extracted from the following subreddits: *askwoman, relationships, relationship_advice, romancestories, retroactive_jealousy, short_stories, sex,* and *teenagers*.

Each of the stories was manually tagged with one of the following three classes: Cheating (*N*=843), Paranoia (*N*=1046), and Other (*N*=940). We used the stories labeled with *Cheating* and *Other* for the cheating vs. no-cheating classification (fine-tuning) task, and the stories labeled with *Paranoia* and *Other* for the paranoia vs. no-paranoia classification (fine-tuning) task.

#### The fine-tuning procedure

The purpose of fine-tuning is to update BERT’s parameters (i.e., the weights of the neural network), as set in the pre-training stage, to ‘point’ toward cheating/paranoia contexts. We used a pre-trained, 12-layers version of BERT^2^ and added a classification head on top of it (the full architecture is detailed in the Supplementary Material and illustrated in Figure S1). For each classification task, we trained this BERT-based classifier (i.e., BERT + classification head) using the backpropagation algorithm (Kelley, 1960) which updates the parameters of both the classification-head and the original BERT model (Figure 2). Importantly, the classification head is removed after training, and the remaining BERT model – that has just been updated – is considered as CheatBERT (in case of fine-tuning with respect to the cheating classification task) or ParaBERT (in case of fine-tuning with respect to the paranoia classification task, Figure 2). All other computational aspects of the entire process are described in detail in the Supplementary Material.

**Figure 2.**
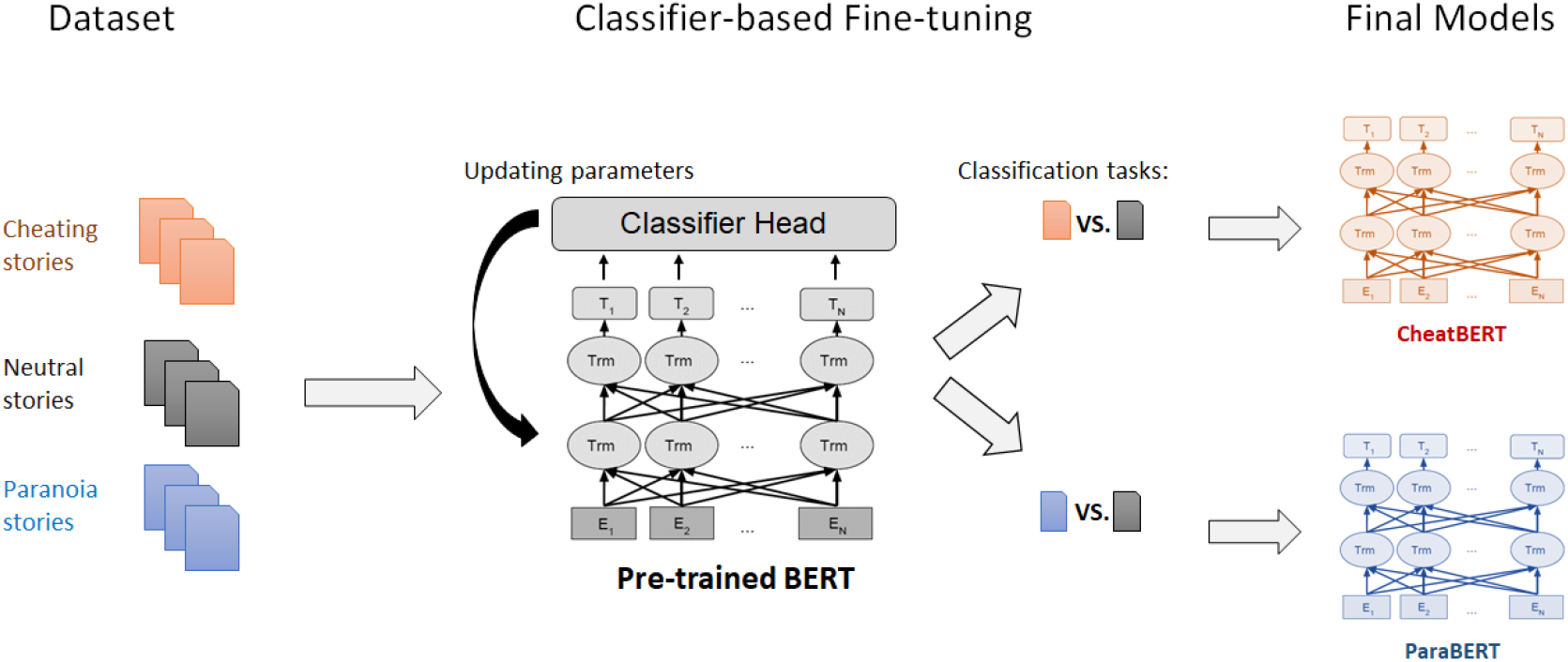
The classifier-based fine-tuning process. We aim to create two new context-dependent language models by changing the parameters of an existing pre-trained model (BERT) to point toward the *cheating* or the *paranoia* context. First, we collected ~3000 short stories, tagged as *cheating, paranoia,* or *neutral* stories. Then, we trained (fine-tuned) a BERT-based classifier (the BERT model with a classification head) on either the cheating vs. no-cheating or the paranoia vs. no-paranoia classification task, updating the parameters of both the pre-trained BERT model and of its classification head. Finally, we removed the classifier head and were left with the new, cheating- (or paranoia-) induced variant of BERT: CheatBERT (or ParaBERT).

#### Control models

Besides CheatBERT and ParaBERT, we created four additional BERT variants using the same fine-tuning procedure, but with different datasets and classification tasks. These models serve as control (baseline) models in our analyses. These variants are divided into two model pairs, where the members of each pair differ from each other in the specific context they model, but their contexts still refer to the same general theme (just as CheatBERT and ParaBERT both refer to the relationships and romance theme). The pairs were SpaceBERT -#x2013; MedBERT (*Med* stands for medicine) and GunsBERT – MideastBERT. The models in the first pair are generally related to *science,* while those of the second pair are related to *politics*.

We used subsets of the publicly available *20Newsgroup* dataset*^2^* for training (fine-tuning) these BERT-based classifiers: SpaceBERT was trained to distinguish texts tagged with *sci.space* (*N*=987) from other-science relevant texts (tagged with *sci*; *N*=991); MedBERT was trained to distinguish texts tagged with *sci.med* (*N*=990) from other-science relevant texts (tagged with *sci*; *N*=991); GunsBERT was trained to distinguish texts tagged with *politics.guns* (*N*=780) from other-politics relevant texts (tagged with *politics*; *N*=685); Finally, MideastBERT was trained to distinguish texts tagged with *politics.mideasst* (*N*=795) from other-politics relevant texts (tagged with *politics; N*=685). Classification results of all models are provided in the Supplementary Material (Table S1).

### Extracting neural and computational representations

The current research examines the relationships between neural and computational representations of the story. Representations were extracted segment-wise in accordance with the above mentioned (in the Behavioral Data section) textual segmentation of Salinger’s story (*N*=179 segments, Figure 1). The BOLD signals of each stimulus-locked voxel (*N*=11783, see in the fMRI Data section) were “down-sampled” from TR resolution (TR=1.5 s) to a segment resolution by averaging all TRs within each segment (Mean number of TRs per segment= 2.51, *SD*=1.63).

We extracted segment-wise computational representations from each of our seven BERT variants (CheatBERT, ParaBERT, the original (pre-trained but not fine-tuned) BERT, and the four control models). The extraction process was identical for all models since they all have the same architecture (12 attention-blocks stacked on top of each other). Each segment was fed to the model, together with a context of additional four segments – the two that preceded and the two that succeeded the relevant segment (whenever possible), as well as with the special tokens: CLS and SEP. A vector representation of the segment was obtained by averaging only the embedding vectors (i.e., the output of layer 12 of the model) of the tokens which belongs to the relevant segment. This procedure yielded, for each model, a 179 (segment) by 768 (the BERT dimensionality) matrix (Figure 1).

Since the dimensionality of the segment embedding vectors (768) is much higher than the number of samples in the data (179), we reduced the dimension of the vectors into 32 using principal component analysis (PCA; see also Goldstein et al., 2021). PCA was calculated separately for each pair of models, and for the original BERT. Reducing to 32 dimensions provides a reasonable balance between the relatively low dimensionality and the relatively large fraction of the original variance preserved after the transformation (71% for CheatBERT-ParaBERT, 80% for SpaceBERT-MedBERT, 72% for GunsBERT-MideastBERT, and 68% for BERT).^3^

### Encoder-based context classification

We aim to show that our fine-tuned models (CheatBERT and ParaBERT) capture the information encoded in the brains of the participants which belong to the corresponding group (CheatBERT for the *cheating* group and ParaBERT for the *paranoia* group). In other words, we hypothesize that the neural signal of each of the subjects would be better correlated with the congruent model (e.g., the CheatBERT model with subjects from the cheating-condition group) than with the incongruent one (e.g., the ParaBERT model with subjects from the cheating-condition, and vice versa). Likewise, according to this hypothesis, we can use the models to *predict* the context in which a given subject listened to the story (cheating/paranoia), by correlating his/her neural signal with both CheatBERT and ParaBERT and checking which model is better correlated (Figure 3a). By this logic, we formulated a voxel-wise classification task through which we can quantify the ‘goodness of fit’ of our models.

**Figure 3.**
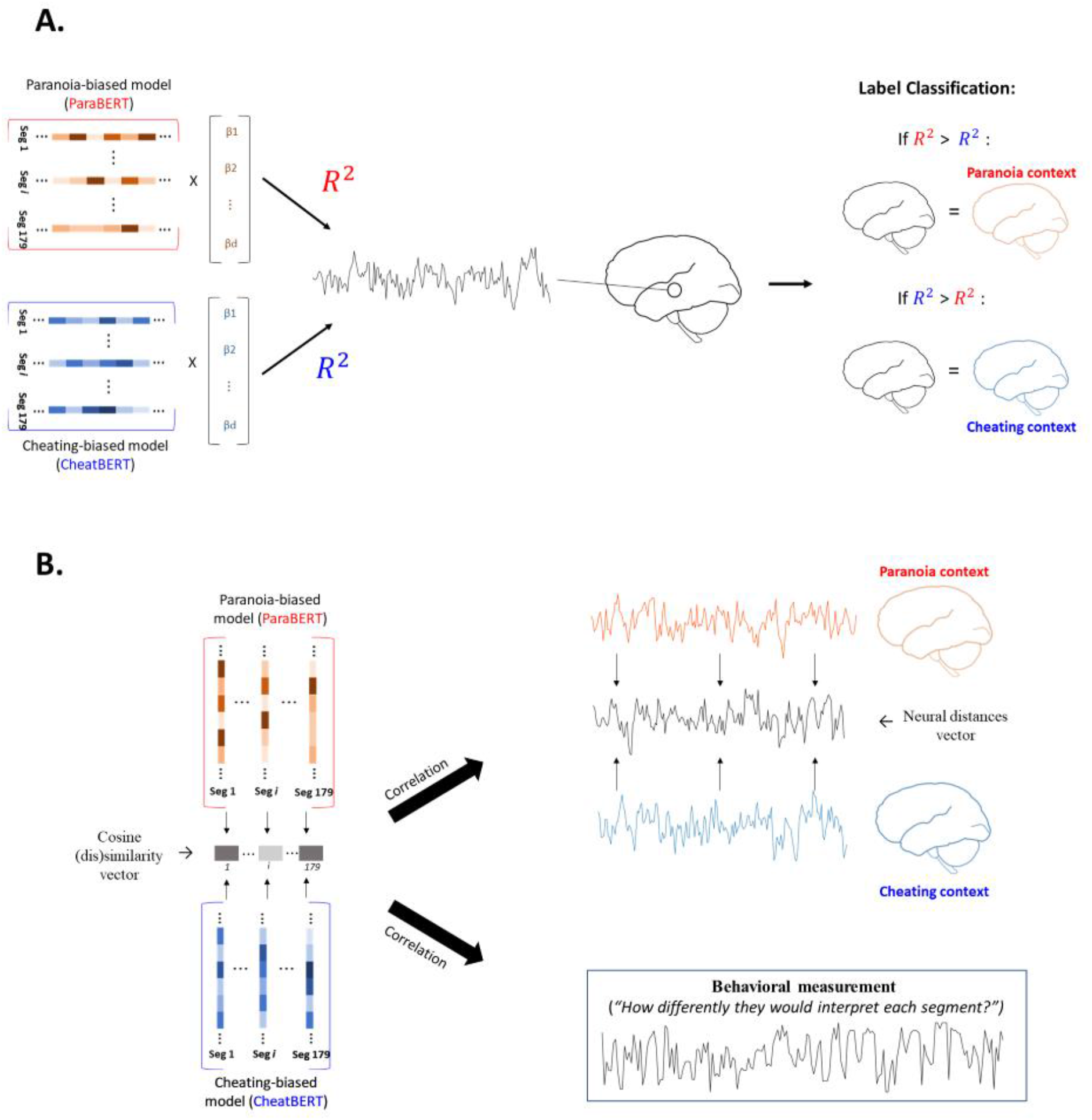
Illustrations of the two primary analyses applied in this paper. (**A**) The voxel-wise encoding-based classifier. The classifier predicts the context in which subjects interpreted the story, by competing the models against each other in their ability to encode the voxel’s BOLD signal. The context would classify as cheating if the CheatBERT-based encoder is better than the ParaBERT-based encoder, and vice versa (measured by comparing the encoders’ 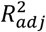 scores). (**B**) The distance analysis. The difference between the cheating-induced and the paranoia-induced interpretations was quantified for each segment, in all modalities: Language models, brains, and behavior. From the language models, we extracted a distance vector by calculating the cosine dissimilarity between models’ vector representations. From the neural data, we extracted a distance vector for each voxel by taking the absolute values of the differences between the averaged signal of the *cheating* group and the averaged signal of the *paranoia* group. The behavioral distances vector was collected by asking five independent raters to rate, for each segment, how differently subjects from different groups would interpret the segment. We analyzed the correlations between the models’ distance vector and the neural distance vectors, and between the models’ distance vector and the behavioral measurement.

For each stimulus-locked voxel (11783 voxels with reliable inter-subject correlation, see in the fMRI Data section), the classifier iterates over all subjects’ brains (*N*=40) and predicts the context (cheating/paranoia) as follows: First, we fit two linear regression models to predict the neural time-course from the vector representations of the story (a.k.a “neural encoder”, see Huth et al., 2019; Goldstein et al., 2021). One model uses vectors extracted from CheatBERT, and the other uses the ParaBERT’s vectors. Then, we calculate the 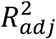 score (adjusted coefficient of determination) from each model and classify the context in accordance with the best 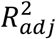 score. Namely, if the CheatBERT’s 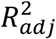 score is higher than the ParaBERT’s 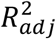 score, we will classify that brain as *cheating,* and vice versa (Figure 3a). We evaluate the classifier by calculating the accuracy rate of its prediction (the number of correct predictions divided by 40). This procedure provides a single accuracy score for each voxel, which quantifies the extent to which our models fit the neural signal in different brain areas.

We repeated the same analysis using other pairs of control models, for the purpose of comparison. The alternative pairs were: CheatBERT vs. BERT, BERT vs. ParaBERT, GunsBERT vs. MideastBERT, and MedBERT vs. SpaceBERT. The significance testing of this analysis is described in the *Statistical analyses* section below.

### Distance analysis

The neural modulation caused by the context is not uniform, but varies throughout the story: there are parts of the story that cause more substantial neural differences between brains compared to other parts of the story (Yeshurun et al., 2017). We wanted to test whether this dynamic is also encoded in our fine-tuned models. To test this, we calculated the distance between each pair of segment embedding vectors (extracted from CheatBERT and ParaBERT) using the cosine-distance (which is equal to 1 minus the cosine similarity of the vectors). This process yielded a 179-dimensional distance vector (corresponding to the 179 segments of the story, Figure 3b). Likewise, we calculated the neural differences (distance) between the average brain activity of the *cheating* group and the average brain activity of the *paranoia* group. This is done by taking the absolute values of the differences between the averaged brain activities of each segment in every stimulus-locked voxel. This process yielded a single 11783 (voxels) by 179 (segments) matrix of neural distance scores. Finally, we calculated the correlation between each voxel’s neural distance and the model’s distance vector using Pearson’s *r* (Figure 3b).

In addition, we analyzed the correlation between the models’ distance vector and the differences in human interpretation of the story (the behavioral measurement, see in the *Behavioral data* section). The analyses were repeated using other distance vectors extracted from the following pairs of control models: CheatBERT vs. BERT, BERT vs. ParaBERT, GunsBERT vs. MideastBERT, and MedBERT vs. SpaceBERT. The significance tests of these analyses are detailed below in the *Statistical analyses* section.

### Statistical analysis

All analyses were tested for statistical significance using non-parametric permutation tests. In the classification analysis (the *Encoder-based context classification* section) we tested the significance of the accuracy scores by creating an estimated null distribution using 1000 permutations of the data. In every permutation step we shuffled the labels (i.e., cheating/paranoia) of the participants, ran the classification analysis on that randomized data, and saved the accuracy scores. The procedure returns a different 1000-sized distribution for each voxel (for a total of 11783 voxels). To account for multiple hypothesis testing, we calculated the *p*-values of the observed (real) accuracy scores using family-wise error rate (FWER; Nichols & Hayasaka, 2003) estimation: We combined the 11783 distributions into a single, 1000-sized distribution by taking only the maximum value (i.e., the best voxel’s accuracy score) from each permutation step. Next, we calculated the *p*(FWER) score from the obtained max-values null-distribution using the following formula:

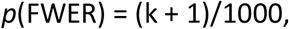

where *k* is the number of max-values larger than the real value. We considered a voxel’s score as significant if its *p*(FWER) was smaller than 0.05.

The same procedure was applied for the remaining analyses and the only difference was regarding the way we permuted the data. In the distance analysis (the *Distance analysis* section) we calculated the *p*(FWER) of each Pearson’s *r* score using the max-values null-distribution obtained from 1000 permutations, as above, but the data was permuted using randomized phase-shuffling. This method randomizes the signal while maintaining the exact mean and autocorrelation as the original signal (Simony et al., 2016). We implemented this shuffling method by applying a fast-Fourier transformation (FFT) on the original signal, randomizing only the phase component of the signal, and then applying an inverse fast-Fourier transformation (IFFT) using the original frequency magnitudes and the randomized phases. The shuffling was performed only on the neural signals. In the models-behavior correlation analysis we shuffled only the behavioral signal.

## Results

### Fine-Tuned Language Models Fit Contextual Modulation in the Brain

We proposed the concept of fine-tuning deep language models as a computational framework for modeling how the states of mind of the listeners modulate their neural responses to the same story. To that end, we fine-tuned (trained) pre-trained deep language models (BERT) to distinguish between cheating and no-cheating stories (CheatBERT) or between paranoia and no-paranoia stories (ParaBERT; Figure 2). Next, we used these models to encode the neural responses in two groups of subjects who listened to the same J.D Salinger story. Each group was primed with a different context (state of mind), of either cheating or paranoia, before listening to the story.

Fine-tuning BERT improved our ability to model context-based unique neural responses of our listeners. First, we used the fine-tuned CheatBERT model to encode the averaged neural responses of listeners who were exposed to the cheating context, and we used the finetuned ParaBERT model to encode the average neural responses of listeners who were exposed to the paranoia context. Both models were highly effective in predicting neural signals of the corresponding group of listeners. The CheatBERT model significantly predicted 8226 voxels (69.8% of all stimulus-locked voxels, 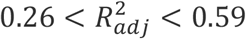, 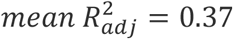) and the ParaBERT model significantly predicted 6692 voxels (56.7%, 0.24 < 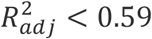, 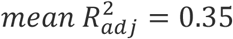; Figure 4a). Next, we analyzed the sensitivity of these models in distinguishing between listeners using our novel encoder-based classification method (Figure 3a). We perform the analysis on a voxel-by-voxel level, among all the stimulus-locked voxels (*N*=11783). The classifier accuracy rate was above the chance level in 921 voxels, ranging from 62.5% to 85% (*p*<0.05, family-wise error corrected). These voxels encompass brain regions in the default mode network (Bilateral TPJ, MFG and Precuneus), as well as language-related areas in the right ventrolateral prefrontal cortex (vLPFC) and bilateral STG and MTG (Figure 4b and Table 1). Hence, neural responses in these brain areas which are unique to each group of listeners (cheating and paranoia group), are significantly associated with the unique information encoded in the corresponding fine-tuned model (CheatBERT and ParaBERT).

**Figure 4.**
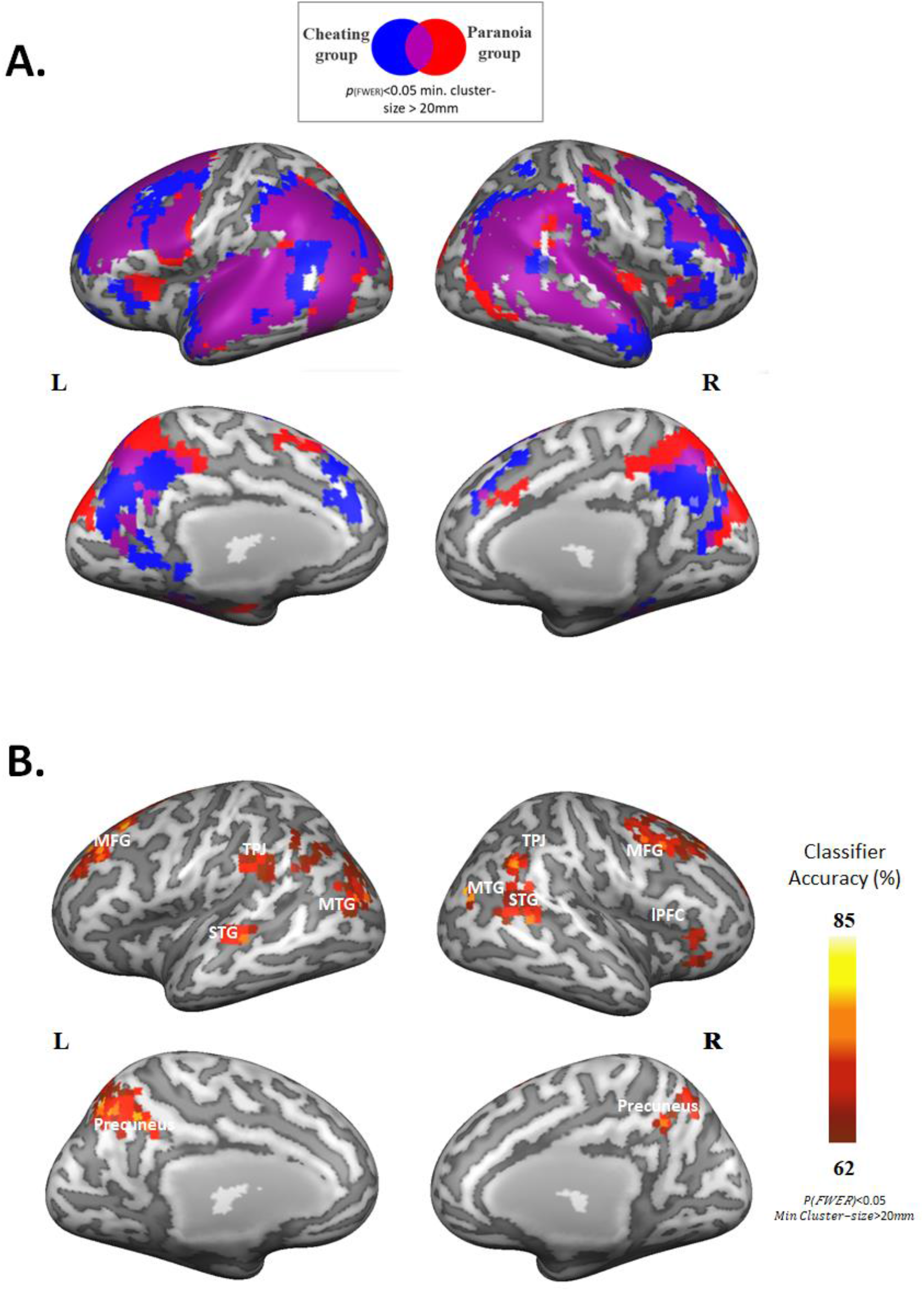
(**A**) Cortical maps showing voxels that are significantly encoded (i.e., with a significant 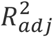 score, *p*(FWER)<0.05, minimum cluster-size>20mm^2^) by the fine-tuned models. Averaged 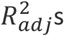 are 0.37 (min-max range: 0.26-0.59) for CheatBERT/cheating group, and 0.35 (min-max range: 0.24-0.59) for ParaBERT/paranoia group. (**B**) The accuracy scores map of the encoder-based classification analysis. The map contains only significant voxels (*p*(FWER)<0.05, minimum cluster-size>20mm^2^). MFG = Middle Frontal Gyrus, TPJ = Temporoparietal Junction, STG = Superior Temporal Gyrus, MTG = Middle Temporal Gyrus, lPFC = Lateral Prefrontal Cortex.

**Table 1.** Brain regions which showed significant classifier accuracy scores.

| Region | Hem. | No. of Voxels | Peak Accuracy score | Coordinates |  |  |
| --- | --- | --- | --- | --- | --- | --- |
|  |  |  |  | X | Y | Z |
| Precuneus | Bilateral | 243 | 0.85 | -6 | -60 | 39 |
| Middle Frontal Gyrus | Right | 120 | 0.725 | 35 | -4 | 41 |
| Middle Frontal Gyrus | Left | 119 | 0.725 | -31 | 15 | 43 |
| Temporoparietal Junction | Right | 69 | 0.725 | 39 | -67 | 35 |
| Temporoparietal Junction | Left | 78 | 0.7 | -42 | -44 | 32 |
| Middle Temporal Gyrus | Right | 15 | 0.775 | 58 | -47 | 10 |
| Middle Temporal Gyrus | Left | 67 | 0.7 | -55 | -52 | 3 |
| Superior Temporal Sulcus | Right | 78 | 0.7 | 53 | -39 | 13 |
| Superior Temporal Gyrus | Left | 70 | 0.7 | -53 | -27 | 8 |
| Ventrolateral Prefrontal Cortex | Right | 62 | 0.65 | 37 | 19 | 11 |

Using the original pre-trained BERT model or the BERT models that are fine-tuned on unrelated contexts did not improve our ability to classify the listener’s state of mind. Running the same classification procedure but replacing ParaBERT with BERT yielded only 25 significant voxels (17 voxels from the vLPFC and 8 voxels from the cuneus, max scores: 70% and 65%, respectively), and replacing CheatBERT with BERT yielded zero significant voxels. Likewise, we found no significant voxels when replacing the ParaBERT and CheatBERT models with the other control-models paires, MedBERT-SpaceBERT and GunsBERT-MideastBERT.

### Models’ distance analysis

The effect of context on the narrative interpretations is not fixed, as some moments in the story are more ambiguous and malleable to shift in context, while others are less open to multiple interpretations. To assess whether our fine-tuned models capture the dynamic fluctuations in interpretability across subjects who listened to the same story while having two opposing perspectives (contexts) we performed two analyses.

In the first analysis, we compared the magnitude of the change in the representation of each segment across the ParaBERT and CheatBERT (cosine similarity vector, Fig. 3B) to the magnitudes of changes in the neural activity, as induced by the paranoia and cheating contexts, in each of the brain areas. Our analysis revealed significant correlations (between *r*=0.3 and *r*=0.43, *p*(FWER)<0.05) in extensive brain areas, including regions from the mentalizing network (bilateral TPJ and Precuneus), the mirror neuron system (bilateral premotor cortex), language-related areas (bilateral STG and right MTG), Bilateral Insula, Right Anterior Cingulate Cortex (ACC) and the left Hippocampus (a total of 1020 voxels, See Table S2 and Figure 5a). Importantly, running the same analysis with other combinations of models (i.e., the pairs: BERT-CheatBERT, BERT-ParaBERT, MedBERT-SpaceBERT, GunsBERT-MideastBERT) did not reveal any significant correlation-maps.^4^

**Figure 5.**
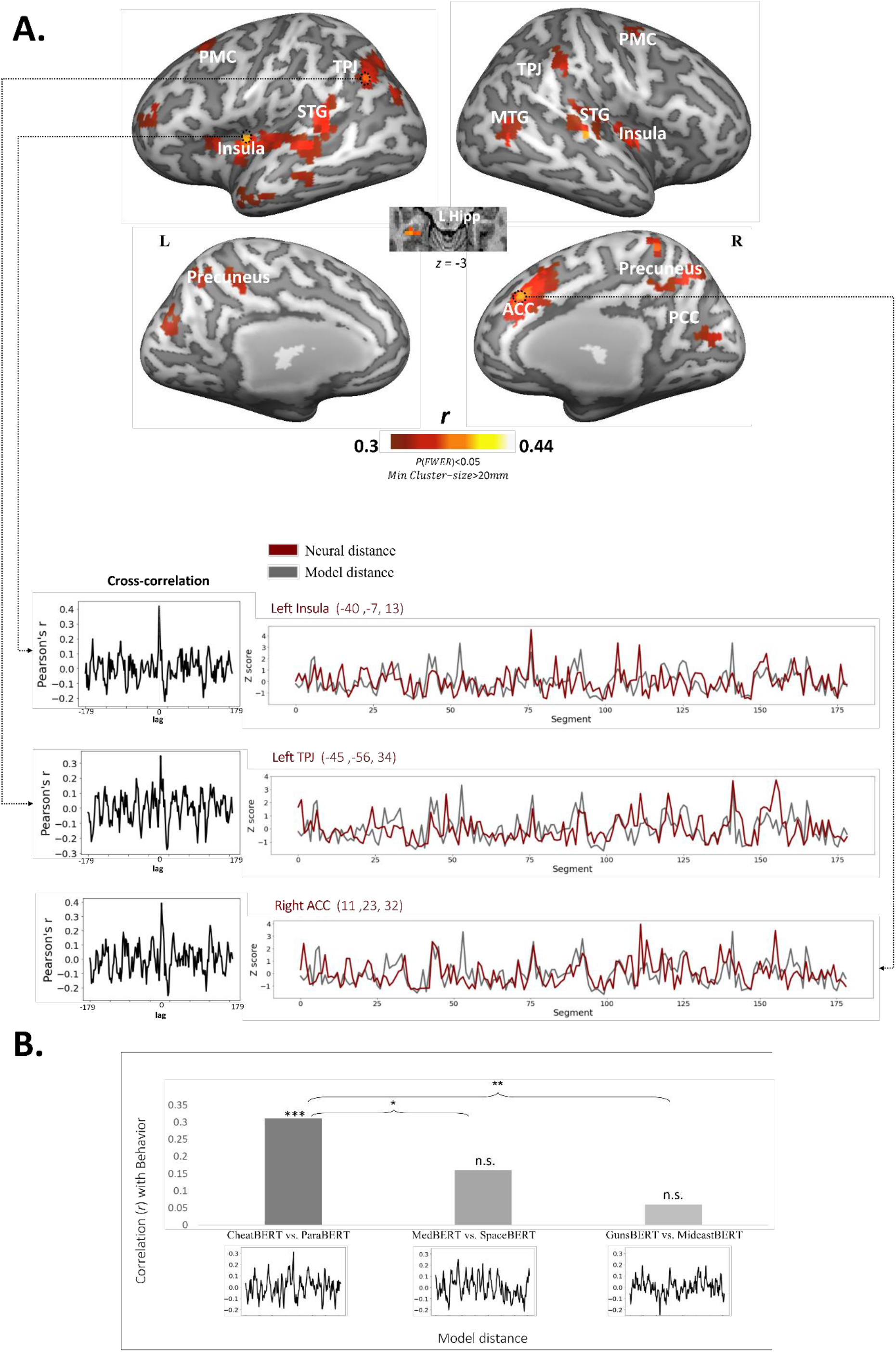
Results of the distance analysis. (**A**) A correlation map showing voxels whose neural distances were significantly correlated with model distance (*p*(FWER)<0.05, minimum cluster-size>20mm^2^). For several brain regions we plot the neural distance fluctuations as measured across segments (maroon colored line), together with the models’ distances (gray colored line). A cross-correlation plot is attached to the plot of each of the regions to visually indicate the signal-to-noise ratio. PMC = premotor cortex, TPJ = temporoparietal junction, STG = superior temporal gyrus, MTG = middle temporal gyrus, PCC = posterior cingulate cortex, ACC = anterior cingulate cortex, Hipp = Hippocampus. (**B**) A bar-plot showing the correlations (Pearson’s *r*) between model distances – as extracted from different pairs of models – and the behavioral scores. Below each bar is the corresponding cross-correlation plot. * = *p*<0.05, ** = *p*<0.01, *** = *p*<0.005, n.s. = non-significant (*p*>0.05).

In the next analysis, we compared the difference in the representation of each segment by ParaBERT and CheatBERT (cosine dissimilarity vector, Fig. 3B) to the estimated change in interpretation between listeners exposed to cheating or the paranoia contexts (behavioral dissimilarity). Behavioral dissimilarity was assessed using independent raters that assessed “how different subjects in the cheating condition and in the paranoia condition would interpret each segment” (see method and Figure 3b). The CheatBERT-ParaBERT distance scores were significantly correlated with the behavioral scores (*r*=0.31, *p*<0.001).In contrast, the correlations between the distances between control models (i.e. the MedBERT-SpaceBERT distance and the GunsBERT-MideastBERT distance) and the behavioral scores were significantly lower (*r*=0.15 and *r*=0.08 for MedBERT-SpaceBERT and GunsBERT-MideastBERT respectively) and not significantly different from zero (p>0.05 for both, Figure 5b).

## Discussion

We presented a computational framework for modeling the effect of listeners’ perspective (state of mind – as defined by their thoughts, beliefs, and prior knowledge) on the way they interpret the exact same textual stimulus. Recent papers found similarities between contextualized word embeddings derived from DLMs (such as BERT or GPT) and the human brain (Mitchell et al., 2008; Huth et al., 2016; Jain & Huth, 2018; Pereira et al., 2018; Gauthier & Levy, 2019; Schwartz et al., 2019; Goldstein et al., 2021, Caucheteux et al., 2021). Building on these findings, we introduced a novel fine-tuning framework to model the effect of listeners’ perspectives on how they process a spoken narrative. Fine-tuning is a method to adjust a pre-trained language model to better fit a narrower linguistic domain. We collected a dedicated dataset (Cheating/Paranoia/Natural stories) which allowed us to fine-tune a well-established DLM, BERT (Davlin et al., 2018), to fit either cheating or paranoia contexts. The process yielded two variants of BERT: CheatBERT and ParaBERT, each induces a different word embedding space (or “semantic space”). Next, we used the two fine-tuned DLMs to model the neural responses of two groups of listeners who listened to the same story after being prompted to have two different perspectives (of cheating versus paranoia).

We consistently showed that the fine-tuned DLMs (CheatBERT and ParaBERT) better fit the neural responses of the subjects with the corresponding perspective (Cheating versus Paranoia). First, we showed that we can use fine-tuned models’ word embeddings to successfully predict the context in which a subject interpreted the story (Figures 3a and 4); Second, we found that the magnitude of change in the representation of each segment between ParaBERT and CheatBERT (measured via cosine dissimilarity) is correlated with the magnitude of change in neural responses between the cheating and the paranoia groups (Figures 3b and 5a); Third, the magnitude of change between the models’ representations of each segment correlated with the perspective-induced difference in interpretation across the two groups (Figure 5b). Importantly, these results are exclusive to the specific pair of fine-tuned models, CheatBERT and ParaBERT, and both models are necessary for modeling the data from the Yeshurun et al. (2017) experiment. Our analyses show that replacing one of the fine-tuned models with the original, pre-trained but not fine-tuned, BERT, as well as using other control contexts to fine-tune the BERT model (as sport and science) yields a substantial reduction in the ability to model listeners’ neural responses as they process the story.

Fine-tuning reshapes (reweights) the relationship among contextual embeddings to better fit the geometrical space of a particular context. For example, the word “wife” might be closer to the word “mine” in the paranoia context, but closer to the word “unfaithful” in the cheating context. The rotation of the geometric space of the language model, as induced by its fine-tuning process, is solely dependent on the difference between the ways individuals use words in the context of cheating and in the context of paranoia, across many situations. Such a fine-tuning procedure is then used as a computational model to model the neural activity of subjects who listened to a new story that was not included in the fine-tuning process. Our results suggest that as subjects change perspectives, they also rotate their semantic space to better represent the relations among words in that particular context. As such, our results provide a new theoretical framework for modeling how subjects’ internal perspectives shape the way they encode incoming narratives to memory.

Interestingly, the human brain has the ability to dynamically reshape its semantic space on the fly, i.e., as listeners process the narratives. In other words, listening to one sentence before the story begins, which provided subjects with either the paranoia or the cheating contexts, was sufficient to alter the geometry of the brain embeddings across subjects in each group. Currently fine-tuning in machine learning is an offline process that requires additional training of the language model. In contrast, fine-tuning of the semantic representations in humans can happen “on the fly” as people encounter everyday stories about cheating and paranoia. It is still not clear, however, how the human brain dynamically shifts among these contexts in real-time. Future work is needed in order to bridge this gap, possibly developing a single model (instead of multiple fine-tuned models) which has the capability of changing its own parameters dynamically when applied in different contexts.

An anatomical investigation of our results implies that the fine-tuning methodology helped the models to encode not just semantic and other linguistic features, but also other high-order characteristics associated with the listener’s state-of-minds, such as their attitudes and their feelings about the characters in the story. Besides several language-related areas (e.g., STG and lPFC), we found models-brain associations in brain areas related to the default mode network (DMN; Mars et al., 2012, see also Yeshurun et al., 2021), such as the TPJ and the Precuneus. These areas were associated with the ability to integrate previous information with the incoming information across longer timescales (i.e., over many minutes; Yeshurun et al., 2020; Friederici, 2020). In addition, the DMN is also linked to the ability to think about other’s mental state, a.k.a. the mentalizing network (Mar, 2011; Saxe & Powell, 2006; Schurz et al., 2014; Spunt & Adolphs, 2014). Moreover, in our second analysis, we found a strong correlation between the distance between the textual representations of the models and the neural dissimilarities between the groups in the ACC and the Insular Cortex (Figure 5a and Table S2). Previous works associated activations in these areas with empathy (Cerniglia et al., 2019), and more specifically with the ability to experience others’ feelings (Novembre et al., 2015; Osaka et al., 2004) and emotions (Singer et al., 2004; Wicker et al., 2003). Therefore, the correlation between our models and neural activities across these functional areas indicates that fine-tuning can alter the geometry of the semantic models along many cognitive dimensions.

From a computational point of view, the current study gives a new perspective on the concept of fine-tuning. Usually, fine-tuning is intended for improving the performance of DLM-based classifiers in downstream tasks. For example, fine-tuning can be used to further improve many downstream tasks such as text summarization, machine translation, sentiment analysis, and so on, by adding a few more training steps that optimize the model’s parameters for the specific task (Devlin et al., 2018). In other words, the fine-tuning stage creates a unique and dedicated variant of the DLM for each downstream task (or “stimulus”). Here, however, we used a different logic: instead of adopting a single model to a stimulus, we use fine-tuning to create multiple models that will later be applied to model the same stimulus (i.e., Salinger’s story) and examine differences in how listener’s with different perspective perceive them. This approach has a great advantage in cognitively motivated computational modeling, since in real life, we may process the same stimulus in different ways, depending on our given state of mind. The present study, therefore, provides the first evidence for the relationship between the way the deep language model of the human brain adapts to changes in perspectives and contexts in the natural world.

## Supplementary Material

### 1. Model architecture and the fine-tuning process

The same architecture was used for both the cheating- and the paranoia-classification models and is illustrated in figure S1. It consists of one classification head located on top of the original pre-trained version of the BERT encoder (‘base’ version, taken from the *Huggingface* repository^5^). Since BERT’s input is limited to a maximum of 512 tokens, and most of our stories are longer, a sliding window method (Pappagari et al., 2019) has been adopted. For each story, a fixed size window was ‘moved’ across the text, with an overlap between the windows (the size of the overlap as well as the size of the windows are both hyper-parameters of the model). Each textual window was fed into BERT, together with the standard prefix and suffix tokens (CLS and SEP), and the output vectors of all its tokens (apart from the CLS and the SEP tokens) were then averaged together, yielding a single window-representation vector.

Next, vectors from all the windows were fed into the classification head, which consists of one attention layer and one fully connected nonlinear layer (with sigmoid activation) on top of it. The final output is a scalar ranging from 0 to 1, which represents the certainty of whether the story is about cheating or paranoia (1) or about other relationships related content (0).

The classifiers were trained to minimize the binary cross-entropy loss, using the batch gradient-descent algorithm (batch-size=4). 70% of the data was assigned for training, 15% for testing, and the remaining 15% for development and hyper-parameters calibration. The hyper-parameters included: window size {128,256,512}, overlap size {0,32,64,128}, the attention layer’s dimension {64,256,512}, the number of neurons in the fully connected layer {32,64,128,256}, and the learning rate {1e-3,1e-4,1e-5}. The training procedure ran epoch by epoch until no improvement was obtained in the model’s predictions on the development data. Model performance was evaluated using a standard accuracy metric on the predicted scores (scores higher than or equal to 0.5 have been considered as 1, and lower than 0.5 as 0), which returns the proportion of the correct predictions.

Importantly, the original BERT’s parameters from the top 3 (out of 12) layers were all updated during the training (the remaining parameters were maintained frozen due to a limited computational power).

**Figure S1.**
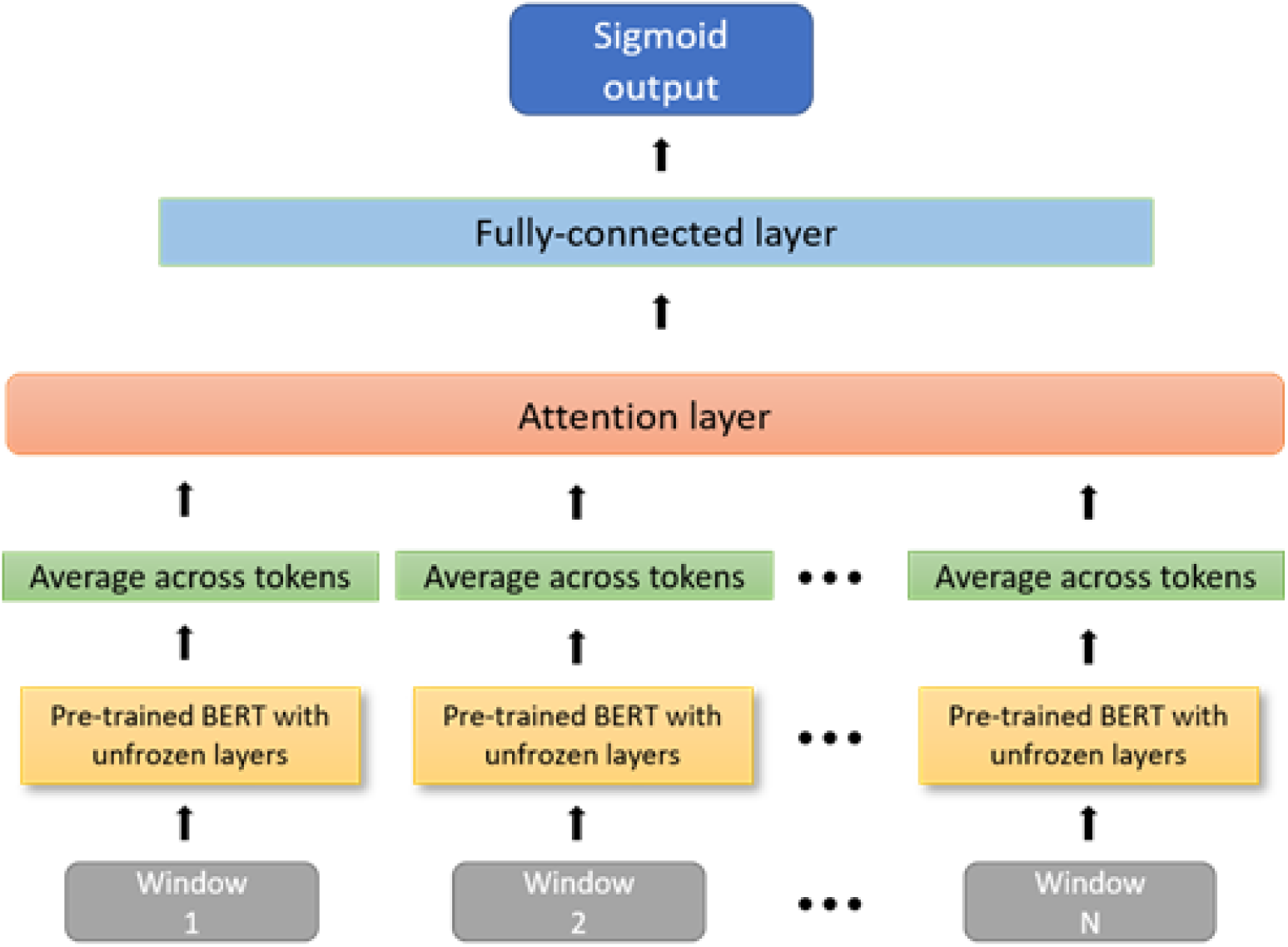
An illustration of the BERT-base classifier.

### 2. Classifier evaluation

We propose a novel classifier-based fine-tuning methodology to produce distinct variants of BERT. The first step in assessing the advantage of this procedure is to evaluate the performance of the difference classifiers. If this method is indeed effective, classifiers should achieve high accuracy scores when tested on the tasks they were trained on, but also, they should show a reduced accuracy in performance on tasks they were not trained on. All six classifiers resulted in a high accuracy score on the test data of their task, ranging from 0.80 to 0.95 (see table S1, the main diagonal). Testing these classifiers on tasks that they were not trained on (e.g. testing the CheatBERT model on the paranoia classification task), indeed leads to a substantial performance drop. This implies that the models are all selectively specialized to the task they were trained on, and each one indeed captures a different and unique context.

Interestingly (and not surprisingly), as can be seen in table S1, the performance declines are not uniform across all novel tasks, but rather they are affected by the global context they are sharing with each model. The performance of the models on novel tasks that belong to their global context (for example, CheatBERT and the paranoia classification task are sharing the same global context, which is *romance and relationships*), are better than their performance on novel tasks that do not relate to the global context (e.g., CheatBERT and the space-classification task).

**Table S1:**
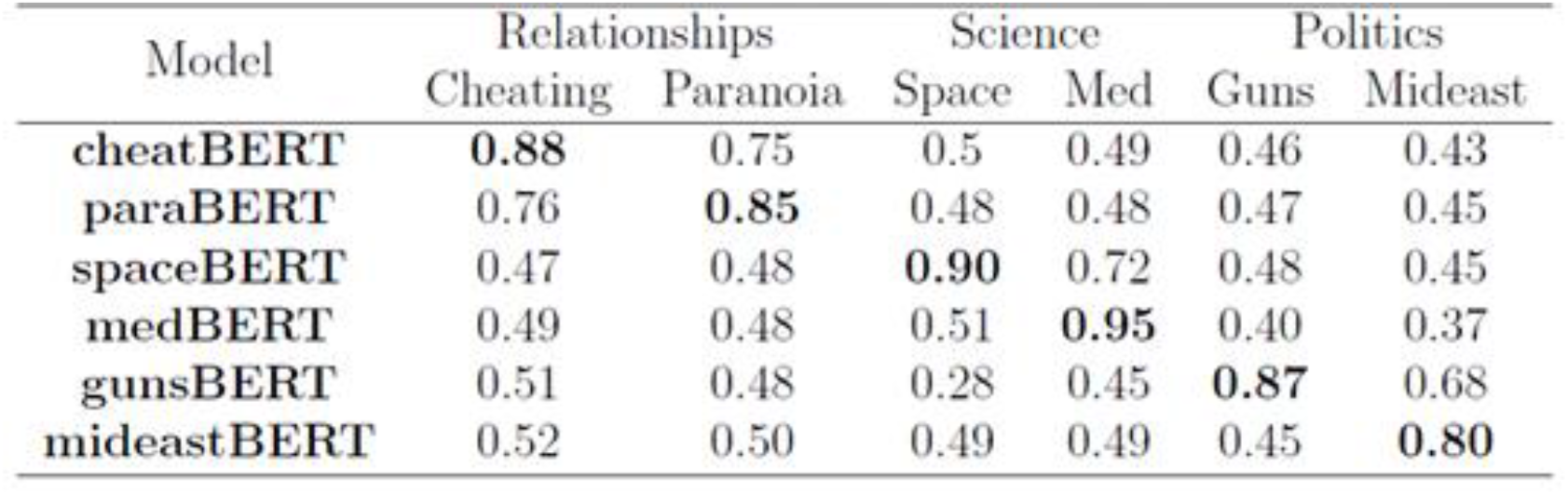
Classifier results by model (rows) and task (columns).

**Figure S2.**
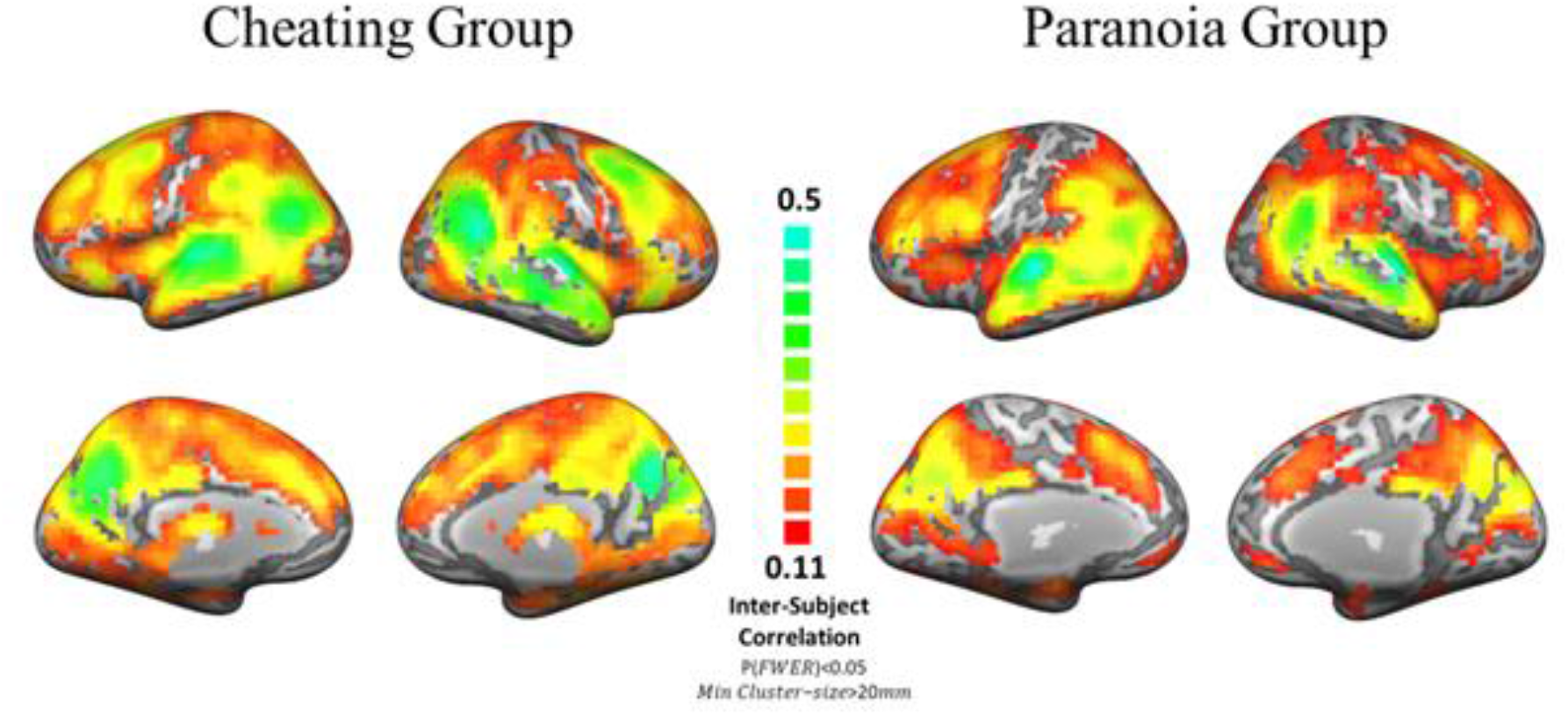
Cortical maps of the inter-subject correlation (ISC, Hasson et al., 2004) scores calculated between subjects within each experimental group.

**Table S2.**
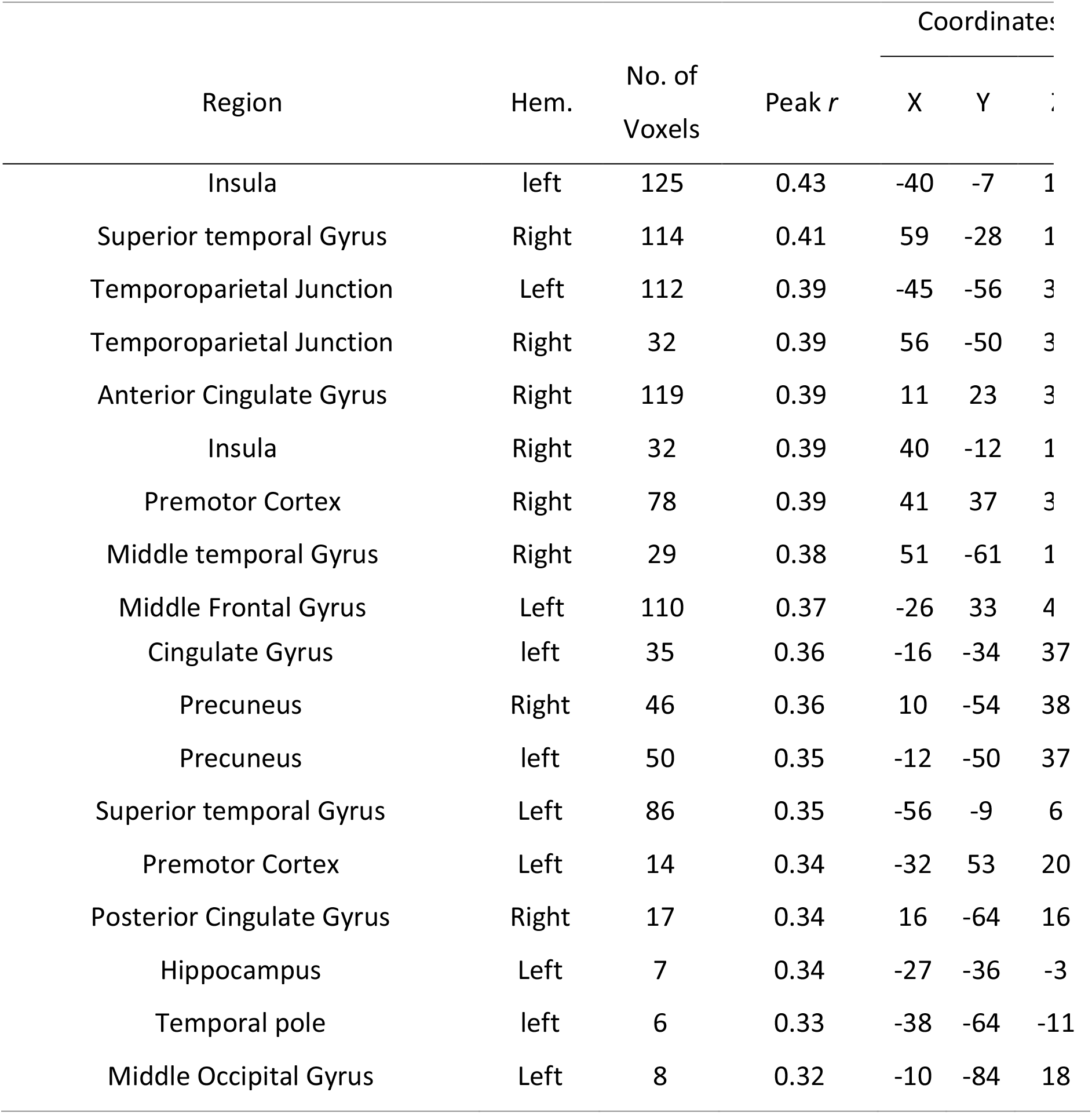
Brain regions whose voxels’ distances were significantly correlated with models’ distances.

## Footnotes

1 https://github.com/google-research/bert

2 http://qwone.com/jason/20Newsgroups/

3 The results below were also replicated using other dimensionalities, ranging from 16 to 75 dimensions.

4 For BERT-ParaBERT, MedBERT-SpaceBERT and GunsBERT-MideastBERT we found zero significant voxels after correcting for multiple comparisons. For BERT-CheatBERT we did find 30 significant voxels, but they did not reach the cluster-size threshold.

5 https://huggingface.co/bert-base-uncased

